# Single cell level analysis of ATP release kinetics and cell fate following ultrasound targeted microbubble cavitation using microscopy techniques

**DOI:** 10.1101/2025.02.02.636151

**Authors:** Marie Amate, Ju Jing Tan, Francis Boudreault, Ryszard Grygorczyk, Thomas Gervais, François T. H. Yu

## Abstract

It is known that ultrasound-targeted microbubble cavitation (UTMC) can induce vasodilation. This image guided spatially targeted approach is called provascular therapy when used as a radiotherapy sensitizer in radiation oncology. Extracellular adenosine-5’-triphosphate (eATP), which plays an important role in vascular tone regulation, is released by cells following UTMC, possibly through sonoporation (formation of temporary and non-deadly pores in the cell membrane) and/or cell death. Herein, we were interested in quantifying UTMC-mediated ATP released *in vitro* using a microfluidics-based model and study its relationship with UTMC-mediated cell fate to better understand and improve UTMC mediated bioeffects.

Lipid microbubbles (MB, Definity®), luciferin-luciferase (LL – for eATP quantification), and propidium iodide (PI – poration tracer) were flown over HUVEC cells cultured in a microfluidic device. Ultrasounds at 1 MHz, varying in pressure (300, 400 kPa) and length (10, 100, 1000 cycles) were applied to the chip. The LL chemiluminescent signal after the ultrasound pulse was acquired with an EMCCD camera to characterize ATP release kinetics. Then, a viability assay was performed with calcein-AM. An in-house MATLAB program pairing eATP kinetics with PI/calcein data was used to classify cells into three categories (sonoporated, dead, and untreated).

Within the testing conditions, a single UTMC pulse caused between 4% and 55% PI-positive (PI+) cells in the ultrasound-treated area. Amongst PI+ cells, we generally found more dead cells than sonoporated cells, except for milder pulses (300 kPa; 10 and 100 cycles). The analysis of individual responses of ATP release demonstrated that dead cells released more ATP (up to 22.4 ± 12.2 fmol/cell) than sonoporated cells (6.8 ± 3.4 fmol/cell) and at a faster release rate which peaked at 4s.

This study showed that sonoporation plays a significant role in UTMC-mediated ATP release, advancing our understanding of UTMC’s potential use as a radiosensitizer in solid tumors.

## Introduction

Tumor hypoxia plays a major role in the resistance to anti-cancer drugs [1] and radiotherapy [2,3]. Indeed, the oxygen level in the tumor is a key parameter for radiotherapy efficacy [2] as it is necessary to stabilize the damage induced by irradiation. Thus, improving perfusion and oxygen supply to the tumor through vasodilation is a promising strategy to improve radiotherapy outcomes [4]. Interestingly, a spatially targeted treatment, called Ultrasound-Targeted Microbubbles Cavitation (UTMC), has been shown to induce vasodilation of the arterial and microcirculation in muscle tissue [5–7] and recently in human prostatic adenocarcinoma [8].

UTMC is an image-guided therapy that can be spatially targeted on tumors by navigating an ultrasound (US) beam. The US pressure excites microbubbles (MB), previously injected intravenously, and circulating systemically in the blood stream. There are several MB formulations clinically used in contrast echography [9]. MB are composed of a biocompatible gas encapsulated in a lipid, protein or polymer shell. In an US field, MB undergo periodic volumetric oscillations, also termed acoustic cavitation. These oscillations can be of small amplitude and regular (stable cavitation) or asymmetrical and chaotic, resulting in MB collapsing and jetting (inertial cavitation). The stable and inertial cavitation of MB can have multiple effects on nearby cells. For instance, UTMC has been shown to induce calcium signaling [10], endocytosis [11], and it has recently been shown to activate the purinergic pathways and the endothelial nitric oxide (NO) synthase (eNOS), known to regulate vascular tone [6,7].

Pioneering work has demonstrated that US therapy alone could initiate a localized provascular response in ischemic tissue in humans [12,13]. The addition of MB was found thereafter to considerably improve the local vasoactive response [5]. The eNOS signaling pathways were shown to be involved not only in the vascular response in the presence of MB [5–7], but also in their absence [14–16]. Indeed, endothelial cells are sensitive to local shear stress, which can lead to NO generation [17,18]. MB cavitation-induced shear stress has been found to be in the kilopascal range [19], far exceeding the level required to activate eNOS, which lies in the range of tens of pascals [18].

The potential role of adenosine-5’-triphosphate (ATP) in the signaling cascade leading to vasodilation has been addressed in prior studies [20]. Extracellular ATP (eATP) activates P2Y receptors (P2Y_1_, P2Y_2_, and P2Y_4_) on endothelial cells and stimulates NO production [21]. Adenosine, the end-product of ATP hydrolysis, can also regulate the vascular tone by stimulating NO production through its action on A_2B_ receptors expressed on endothelial cells [22,23], or on A_2A_ receptors on smooth muscle cells [20,21]. Thus, eATP is a critical molecule regulating vascular tone, and *in vitro* investigation demonstrated that it could be released via lytic and non-lytic pathways using UTMC [6,24,25]. However, the individual kinetics of release and contribution to ATP secretion from these UTMC-induced pathways are largely unknown.

One interesting phenomenon induced by UTMC is sonoporation. Sonoporation refers to the formation of temporary pores in the cell membrane by sonication, followed by its resealing, which is necessary for cell survival. Pore formation is possibly caused by the shear stress augmentation on the cell membranes, and liquid-jet formation in the MB collapsing during inertial cavitation [26,27]. These cells will be designated as “sonoporated cells” in this study. Pore size varies from tens to hundreds of nanometers in diameter [11,28], and the pore’s resealing process starts a few seconds after opening and lasts a few tens of seconds to 1 min [28]. A pore remaining open for more than 2 min, however, is likely to cause cell death [29,30]. Such a cell will be referred to as a “dead cell”.

Pore formation, followed or not by cell death, in endothelial and red blood cells is a source of lytically released ATP [6,24,25]. This eATP could also be secreted by non-lytic mechanisms via the activation of the vesicular pathway of ATP, triggered by shear stress on endothelial cells [23] and/or the opening of conductive ATP channels. Although the role of pannexin-1 channels in ATP release remains controversial [25], they have been suggested to contribute to eATP detected in sonicated tissue up to 30 min following UTMC [6,31].

The release of ATP by sonoporated cells is particularly interesting. Even though sonoporated cells remain alive for a short time after sonoporation [29,32], they could form a large reservoir of ATP, contributing to the provascular response [17]. Accordingly, a relatively high quantity of ATP has been shown to be released by cell layers containing sonoporated cells combined with low UTMC-induced mortality [6]. To our knowledge, the direct release of ATP from sonoporated cells has of yet neither been observed nor quantified *in situ*.

In this study, we explored the effect of UTMC on the ATP release kinetics from dead and sonoporated cells in a microfluidic model. We developed a fluorescent microscopy method for the detection of sonoporation and cell death, and adapted a real-time imaging approach to quantify eATP [33] in our microfluidic device. Together, these methods enabled us to characterize the kinetics of ATP release from pores created by UTMC.

## Methods

### Microfluidic device design and manufacturing

The microfluidic device was designed in-house with the software SolidWorks® (Dassault Systèmes, France). The design consisted of three square-section straight parallel channels (600 μm × 600 μm × 2 cm) spaced by 800 μm, designed to fit within the −6 dB amplitude at the transducer’s focal point (Fig 1A). The channels were reunited at the extremities to have one inlet and one outlet. The entire volume of the chip was 42 μL. The volume used for the injection of any reagent was 50 μL and the washing volume used for flushing was 100 μL.

**Fig 1.**
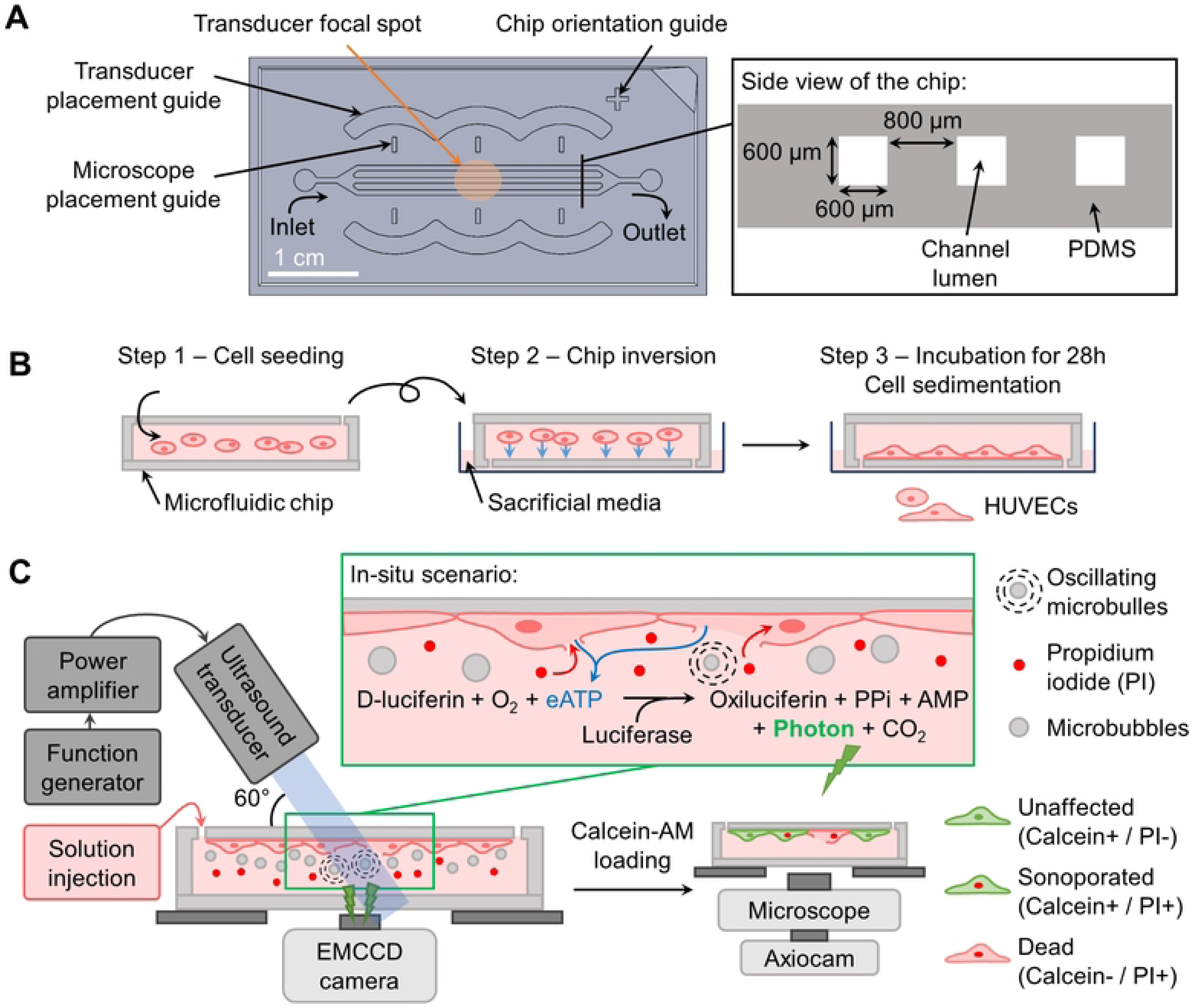
Experimental methods. (A) Description of the microfluidic design. (B) Schematic of the cell culture method (side views). (C) Schematic of the experimental setup and imaging principles (side views). (PPi: inorganic pyrophosphatase; AMP: adenosine monophosphate).

The manufacturing process of the devices is detailed in the Supporting Information (S2 Appendix).

### Ultrasound setup

The ultrasonic transducer (A303S-SU, 1MHz, 0.5 inch, Evident, Olympus, Waltham, MA, USA) was positioned at a 2.67 cm distance from the surface of the microfluidic device with an angle of 60° with the horizontal axis. The microscope and the US beam were co-aligned at the same focal spot, considering the US refraction angle, using an in-house-designed 3D printed device. The transducer was driven by a function generator (33500B, Keysight Technologies, Santa Rosa, CA, USA) connected to a radiofrequency power amplifier (55 dB nominal gain; Electronics & Innovation 1040L RF Amplifier, Rochester, NY, USA). The US treatment consisted of a single sinusoidal pulse, at a frequency of 1 MHz, triggered manually. The pressure tested were 300 and 400 kPa with different cycle lengths (10, 100, and 1000 cycles). The calibration of the transducer is described in the SI (S1 Appendix).

### Cell culture

Human umbilical vein endothelial cells (HUVEC) (200p-05n, Cell Applications, Inc., San Diego, CA, USA) were sub-cultured in a reconstituted human endothelial cell growth medium (213-500, Cell Applications, Inc., San Diego, CA, USA) supplemented with 1% of penicillin-streptomycin (450-201-EL, Wisent Inc., Saint-Jean-Baptiste, QC, Canada). Cells were incubated in a CO_2_ incubator (5% CO_2_; 37°C) between passages. The passages were done at a confluence not exceeding 80%, and cells were used until the 7^th^ passage.

Before the experiment, cells were washed with Hank’s balanced salt solution (HBSS) (311-512-CL, Wisent Inc.) and harvested using a Trypsin/EDTA solution (325-043-EL, Wisent Inc.) diluted 10 times in HBSS (final concentration of 0.025% Trypsin; 0.221 mM EDTA). The trypsin activity was neutralized with fresh culture medium, and cells were centrifuged (200×g for 6 min) (2150, Megafuge, Baxter, Germany). Cells were suspended in fresh medium at a concentration of 2.5 × 10^6^ cells/mL.

Prior to cell seeding, the microfluidic devices were autoclaved for sterilization and the outer part of the chips was washed with ethanol 70%. The inner part of the chips was washed 3 times with ethanol 70% and 3 times with phosphate buffer saline (PBS) (10010-023, Thermofisher Gibco). Chip channels were coated with fibronectin bovine plasma (F1141, Sigma-Aldrich) at a concentration of 100 µg/mL in PBS for 1 h at room temperature. Chips were washed 3 times with fresh medium and then 100 μL of the HUVEC cell suspension (2.5×10^5^ cells/mL) was injected slowly in the chips (Fig 1B – Step 1). The chips were immediately flipped with the outlets facing the bottom of a Petri dish (diameter: 6 mm) previously filed with 2 mL of sacrificial medium (Fig 1B – Step 2). The chips were incubated (5% CO_2_: 37°C) for 24 h before the experiment and the medium was changed at 8 h and 22 h after cell seeding.

### Imaging setup

The chemiluminescence signal of eATP was captured in real time with the setup described by Tan *et al.* [33]. An electron-multiplying charge-coupled device (EMCCD) camera (512 × 512, 13-µm pixel size, camera Evolve 512, Teledyne Photometrics, Tucson, AZ, USA) was placed under the mechanical stage supporting the microfluidic device. A standard c-mount lens was mounted on the EMCCD camera objective, to achieve a magnification of 0.33×, allowing a field of view (FOV) of 20 mm × 20 mm (256 × 256 pixels per image). All the acquisitions were done at 1 frame/s and an integration time of 1 s. To maximize the signal, a binning of 2 × 2 was applied allowing a final resolution of 78 μm × 78 μm per pixel.

The acquisition of the fluorescent images was done with an inverted epifluorescence microscope (Axio Observer Z1, Carl Zeiss MicroImaging, Oberkochen, Germany) with an automated 2-axis stage able to scan the surface of the chip. The calcein and the propidium iodide fluorophores were imaged with a filter set 62 HE (Carl Zeiss MicroImaging) modified with a 530/50 emission filter (Chroma Technology Corp, Bellows Falls, VT, USA) to reduce signal cross-talk, and a filter set 02 (Carl Zeiss MicroImaging) respectively. Cells were illuminated for 1 s at 505 nm for the acquisition of the PI signal and 200 ms at 470 nm for the acquisition of the calcein signal, with a Colibri light source (Carl Zeiss MicroImaging). A total of 80 images per chip (20 × 4 images, magnification 5×) were captured (AxioCam, Carl Zeiss MicroImaging) to cover a total area of 20 mm × 4.4 mm of the chip. A 20% overlap between images allowed a consistent image stitching of the entire scanned area using the “Grid/Collection stitching” plugin in Fiji [34]. The fluorescence images were also taken with a binning of 2 × 2 and have a final resolution of 2.58 μm × 2.58 μm per pixel. The AxioCam, the AxioObserver automated stage, and the EMCCD camera were controlled numerically via the AxioVision software (AxioVs40 V 4.8.2.0, Carl Zeiss MicroImaging GmbH, Jena, Germany).

### eATP measurement method

#### Enzyme preparation

The eATP was measured with a chemiluminescence luciferin-luciferase (LL) assay (Fig 1C). LL mix powder (Adenosine 5’-triphosphate assay mix; FLAAM-5VL, Millipore Sigma) was reconstituted with cold sterile double-distilled water to form the LL solution. This solution was incubated for 1 h on ice to ensure the total dissolution of the powder in water, and then aliquoted and stored at −20°C for up to 3 months. Aliquots were not reused after thawing to ensure reproducibility. Before eATP measurements, the LL solution was isotonically adjusted (4.5:1 v/v) with a 5-time concentrated physiological solution (in mM: NaCl, 700; KCl, 25; HEPES, 50; (MgCl_2_ · 6H_2_O), 5; (CaCl_2_ · 2H_2_O), 5; D-Glucose, 50).

#### Activity test

The enzyme activity of the total amount of LL solution used within an experiment day was measured at the beginning of the day, by mixing at equal volumes (25 µL each) the isotonic LL solution with an ATP solution diluted in Phenol red-free RPMI (350-046-CL, Wisent Inc.) (ATP final concentration: 100 nM). The mix was briefly pipetted up/down 3 times, and then light emission was immediately recorded in a luminometer (TD 20/20 luminometer, Turner BioSystems, Sunnyvale, CA) with a 10-second integration time. The activity ratio (AR) was obtained from the known ATP quantity (mol) divided by the integrated luminescence signal in the light unit (LU).

#### eATP quantification

We used the imaging-based wide field-of-view quantification method described by Tan *et al.* [33], with some adaptations to accommodate our microfluidic device. As the EMCCD detector is flat, the chemiluminescence signal captured is a projection of the emitted photons in the volume of the chip’s channels. A calibration factor (CF) specific to our microfluidic device was determined to quantify eATP.

The isotonic LL solution was mixed with an equal volume (25 µL each) of ATP solutions diluted in Phenol red-free RPMI at different ATP concentrations (0.1, 0.5, 1, 5 and 10 µM). The mixed solution was immediately injected inside the cell-free microfluidic device and the chemiluminescence signal was acquired for 1 min. The device was washed 3 times with HBSS between each acquisition. The maximum chemiluminescence signal (1-second integrating time), was used to calculate CF as follows:

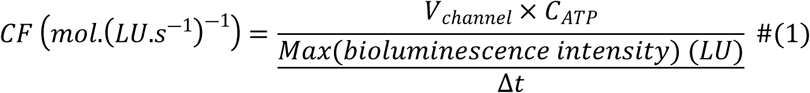

With *V_channel_* (L), the channel volume within the fixed EMCCD camera FOV (*V_channel_ =* 2 cm × 600 µm × 600 µm = 7.2 × 10^-7^ L), *C_ATP_* (mol·L^-1^), the known final ATP concentration inside the chip, and Δ*t* (s) the integration time (1 s). CF was calculated separately for each channel and each concentration. The average of 15 measures gave a CF of 4.6 ± 0.3 amol(ATP)/LU/s (see S3 Appendix).

CF determination was done once at the beginning of this study. To ensure the reproducibility of the measures and minimize the inherent day-to-day variability of the enzymatic LL reaction, we verified the enzyme activity on each day of the experiment as described in the *Activity test* section (see above). An activity correction factor (ACF), defined by the AR measured on the day of calibration over the AR measured on the day of the experiment, was used to compensate for any deviation.

The ATP quantity present inside the chip during an acquisition was determined with this formula:

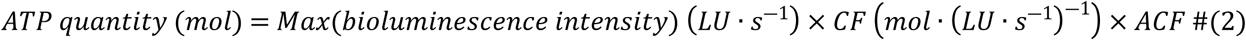

### Experimental design

At the start of the experiment, the cell culture chips were removed from the incubator and flipped back in the original position, making the outlet accessible. The cell layer was consequently lining the ceiling of the channels, maximizing MB/cell interaction by buoyancy (Fig 1B – Step 3). To assess cell viability before the experiments, a propidium iodide (PI, P1304MP, Thermo Fisher Scientific, Eugene, OR, USA) solution (25 µg/mL diluted in PBS) was injected inside the chips and a first scan of PI was done. Viability of over 98% had to be observed to proceed with the experiment.

The isotonic LL solution was mixed at equal volume with a MB (Definity™, Lantheus, Billerica, MA, USA) solution (final concentration: 10^7^ MB/mL, MB/cell ratio of 20) diluted in phenol red-free RPMI medium, also containing PI (final concentration: 25 µg/mL). This solution LL+MB+PI was injected inside the chips. The US pulse was given 20 seconds after the onset of luminescence signal acquisition lasting 5 min in total. In the following analysis and figures, the pulse is considered to be given at time zero second.

After the end of the luminescence signal acquisition, a calcein-AM (C3100MP, Thermo Fisher Scientific, Eugene, OR, USA) solution (4 µg/mL diluted in HBSS) without PI was injected inside the chip. After calcein-AM loading (30 min; 5% CO_2_; 37°C), a scan of the calcein and PI was performed to allow image stitching and cell poration classification (Fig 1C).

### Fluorescent image processing for cell poration classification

Fluorescence images were analyzed with an in-house MATLAB (version 9.11.0 (R2021b) for Windows, MathWorks Inc., Natick, MA, USA) program designed to count cells and detect sonoporation. Cells were sorted into three categories: dead, unaffected, sonoporated cells. The program is further described in the SI (Section S4).

The analysis of the cell proportions in the three categories was done by placing an elliptic region of interest (ROI) on the eATP signal images and the corresponding fluorescent images (Fig 2A). The ROI had a minor axis of 3.5 mm fitting the three channels, in the dimension perpendicular to the chip’s channels direction, and a major axis of length 3.5 mm / sin(60°) = 4 mm, parallel to the chip’s channels direction.

**Fig 2.**
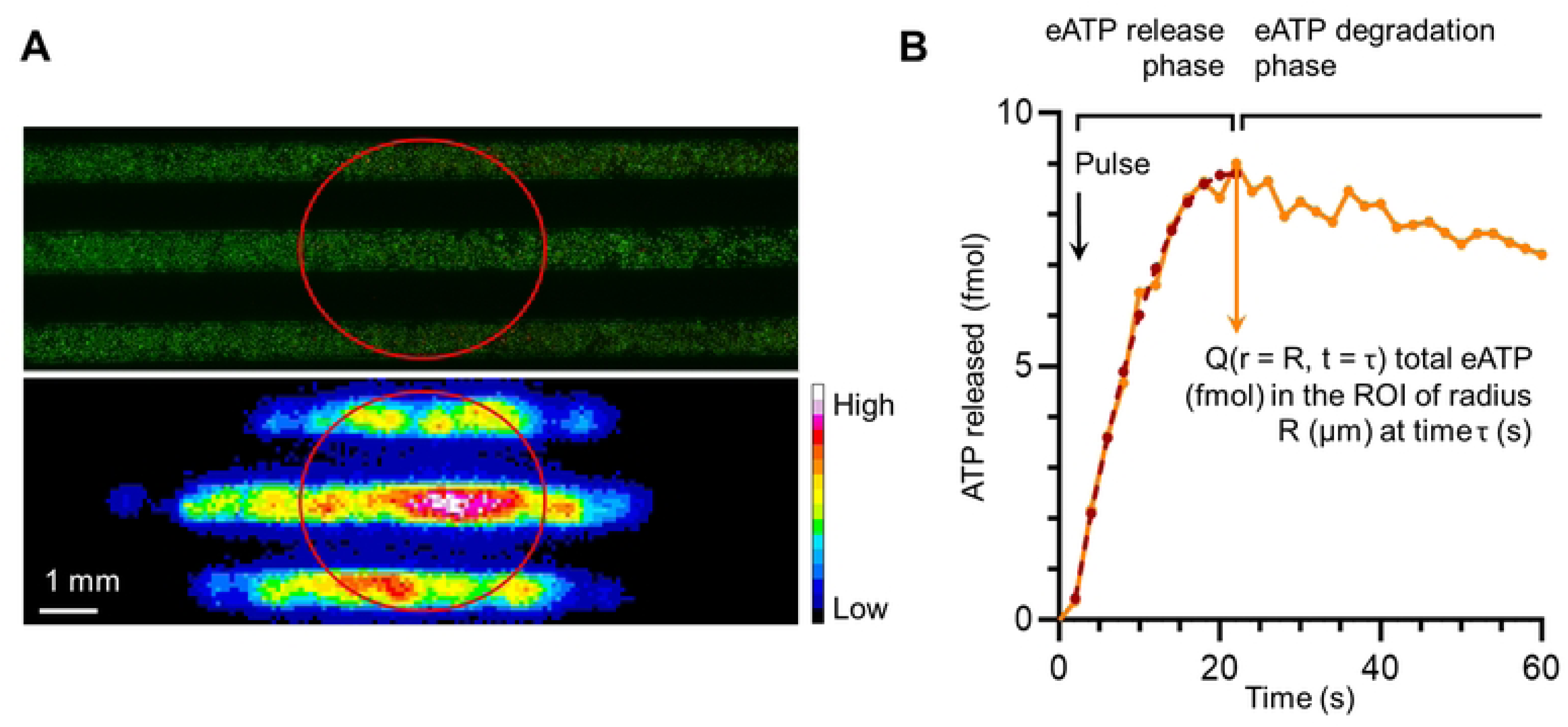
ATP release analysis. (A) Top. Calcein/PI fluorescence image of a chip. Bottom. Bioluminescence image of the same chip treated with ultrasound (300 kPa; 10 cycles). On both images, ROI_US_beam is drawn. (B) Example of an eATP release kinetic profile for an isolated cell in ROI_3.5px. Orange line: experimental data; dark red dotted line: polynomial model corresponding.

Cells present on the entire images (dimension: 7,680 × 1,357 pixels with the Axiocam camera) and inside the ROI_US_beam (dimension: 1,550 × 1,357 pixels) were counted and classified. Chips for which the cell density was below 150 cells/mm^2^ (N = 6) or above 450 cells/mm^2^ (N = 1) were excluded from the study. A total of 36 chips with an average density of 300 ± 71 cells/mm^2^ were analyzed, 16 chips treated with 300 kPa (10 cycles: 6, 100 cycles: 4, 1000 cycles: 6 chips) and 20 treated with 400 kPa (10 cycles: 6, 100 cycles: 8, 1000 cycles: 6 chips).

The sonoporated and dead cell proportions were defined respectively by the sonoporated and the dead cell count divided by the total number of the cells present in ROI_US_beam. Moreover, the gap between dead cells and sonoporated cells ((dead cell count – sonoporated cell count) / PI+ cell count) was calculated to identify the chips where there was more cell death than sonoporation among the PI+ cells.

### eATP images processing

#### Global eATP signal analysis

The ROI_US_beam was drawn on the eATP signal images (Fig 2A) and the total luminescence in the ROI was measured. The luminescence signal inside ROI_US_beam was normalized by the total number of cells in the ROI and by the number of PI+ cells in the ROI.

#### Different eATP signal patterns

Two main luminescence signal patterns were observed on the eATP images: (1) isolated cells, for which the eATP signal remained spatially confined around the cell in the entire cineloop (Fig 3A); and (2) responding-cell clusters, where the signal of several cells was merging at a certain time point in the sequence (Fig 3B & 3C). Different quantification strategies were thus used and compared to quantify eATP.

**Fig 3.**
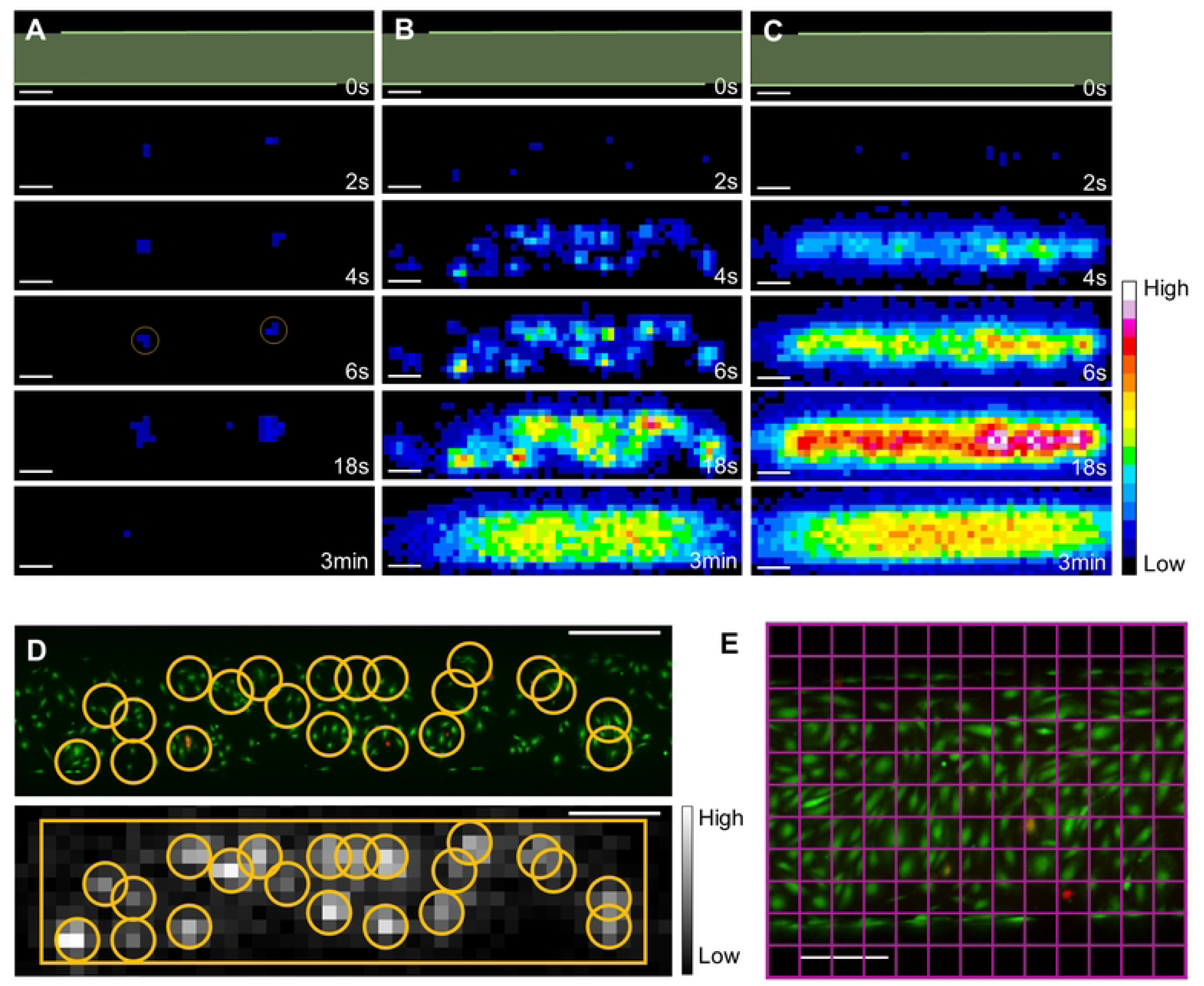
ATP release patterns and isolated cell analysis. (A) Images of the eATP signal over time treated with 300 kPa 10 cycles. We can see two isolated cells responding to the treatment (yellow circles). (B & C) Images of the ATP released by a cluster of responding cells over time treated with 300 kPa 10 cycles and 400 kPa 10 cycles, respectively. In figure A, B and C, the greenish bands on the images at time 0 s show the channel area. Scale bars: 500 µm. (D) Calcein/PI fluorescence image of a cell cluster (top), and the corresponding eATP signal image (time 4 s) with the ROI_1.5px drawn inside the cluster. The rectangular ROI, called ROI_cluster, was used to select the area for the global ATP signal. Scale bars: 200 µm. (E) Calcein/PI fluorescence image of a channel with the drawing of a grid (in pink) representing the pixel matrix of an eATP signal image. Scale bars: 200 µm.

#### ROI sizes determination for isolated cell analysis

eATP luminescence was measured in a static flow condition and the heat in the chips was assumed to be homogeneous. Thus, we assumed that molecule transport was dominated by diffusion. The model of Brownian motion of particles in three dimensions [35–37], was used to estimate the displacement of eATP molecules over time after the pulse. The Mean Square Displacement (MSD) was calculated to assess the diffusion distance of eATP molecules:

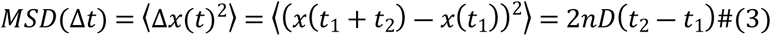

With *n* the dimension of the problem, *D* the diffusion coefficient. As the diffusion of eATP is in three dimensions *n* = 3.

We considered the total amount of the eATP molecules to diffuse in a semi-finite isotropic medium with a constant diffusion coefficient, as it is constrained to diffuse above the cell surface only. The diffusion coefficient for ATP depends on pH, temperature, and the ions present in the solution with the latest estimation at 25°C of 3.68 ± 0.14 × 10^-6^ cm^2^.s^-1^ [38].

#### ATP release by isolated cells

We used a circular ROI of radius 3.5 pixels (273 µm; named ROI_3.5px) centered on the luminescence signal (Fig 3A) which encompassed the entire signal during the whole cineloop. A concentric ROI of radius 1.5 pixels (117 µm; named ROI_1.5px) was drawn. The information on the cell category was determined by the classification program.

For every ROI, the pixel values were summed and converted into eATP quantity using the calibration method described earlier (see “eATP measurement method” section). The background signal was removed by subtracting the average signal from three images before UTMC (20, 18, and 16 s before the pulse was given). A typical curve profile of the luminescence signal is plotted in Fig 2B. We can observe two kinetic phases in the profile; the first kinetic phase, where the signal increases until a peak, and the second kinetic phase, where the signal decreases slowly. The peak of the profile is considered the total ATP released inside the ROI [33], and the time between the pulse and the peak of release was denoted τ (s). When the ROI is a disk of radius R (pixel), the quantity of eATP at a time T (s) is noted *Q(r = R, t = T)*. The release rate ((fmol/s)/cell), was defined by the difference of the eATP signal between two frames divided by the time interval (i.e. 2 s).

##### ATP release model

The model of release during the first kinetic phase for every isolated cell was done with an in-house MATLAB program using the function lsqcurvefit (nonlinear curve-fitting problems in least-squares sense). The most appropriate model was a semi-parabola (Fig 2B) with the vertex (*t_vertex_; q_vertex_*):

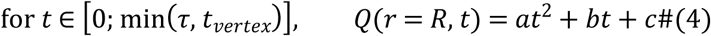

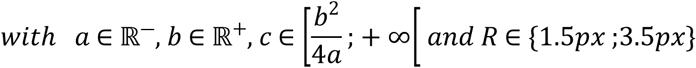

The model was computed for each ROI size (1.5 pixels and 3.5 pixels) for every isolated cell. This gave two sets of parameters for every isolated cell. Each model was normalized by dividing *Q(r = R, t)* by *q_vertex_* at every time point.

#### ATP release by responding-cell clusters

Given the promiscuity of the cells inside a responding-cell cluster, the ROI radius size needed to be optimized. As later shown in the results, the 4-s time point drew our attention. The random walk of diffusion (eq. 3) applied in the three dimensions indicates that the MSD of a molecule of ATP released 4 s after the pulse is 94 µm. According to this calculation, at 4 s after the pulse, all the ATP released should encompass an ROI of radius 1.5 pixels (94 µm < 117 µm).

eATP signals in responding-cell clusters were processed (1) globally and (2) individually: (1) In the global processing, we drew a rectangular ROI of height 11 pixels (named ROI_cluster) and a width variable according to the length of the cluster (Fig 3D); (2) In the individual analysis, we placed a circular ROI of radius 1.5 pixels (ROI_1.5px), in the center of the eATP signal to capture the early moment of ATP release before the signal merged with that of neighboring cells (Fig 3D). The punctual signals identified in the individual analysis were believed to be the ATP released by a membrane poration or disruption. Nevertheless, the literature suggested a release by other non-lytic pathways [6,31], which could be not identified by a punctual signal and a more homogeneous release along the sonicated area. To attempt verification of this hypothesis, we wanted to study the relationship between the sum of the individual signals detected inside ROI_cluster and the total signal integrated into ROI_cluster with two approaches: the detailed sum and the weighted sum approach.

##### Detailed sum approach

We wanted to see if the eATP measurement estimated on the 4-s frame could predict the ATP released inside ROI_cluster. In the case of an ATP released exclusively by sonoporated or dead cells, we can define *Q_DS_* as the total ATP quantity released by cells by:

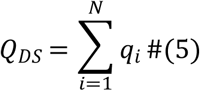

With *q_i_* the quantity of ATP released by the cell *i*, and *i* going from 1 to N the total number of cells inside the responding-cell cluster.

##### Weighted sum approach

Alternatively, we also used the mean ATP quantity released from isolated cell experiments to estimate the global ATP release. In this approach, we can define *Q_WS_* the total ATP quantity released by cells in the cluster by:

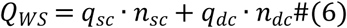

With *q_sc_* and *q_dc_* the average quantity of ATP released by sonoporated and dead cells, respectively (values from the isolated cell experiments), and *n_sc_* and *n_dc_* the number of sonoporated and dead cells respectively.

### Impact of cell confluence on ATP release

The impact of the cell density (cells/mm^2^) on the sonoporation and the ATP release was investigated (Fig 9A). One group of 8 chips (referred to as the “low cell density” or LCD group) was added to the study. The LCD group (84 ± 25 cells/mm^2^) was treated with a single 300 kPa / 10 cycles pulse. This group was compared with the 11 chips (referred to as the “high cell density” or HCD group) described earlier, treated with the 300 kPa / 10 cycles pulse and presenting a full confluence (296 ± 48 cells/mm^2^) (Fig 9B). The cell surface of 107 cells (52 cells in four chips in the group LCD and 55 cells in four chips in the group HCD) was measured to assess the concentration of ATP in cells at the two cell densities. The cell volume was modeled as a prism of height 5 μm and of base corresponding to the cell surface.

### Statistical analysis and data presentation

The statistical analysis and the graph representations were performed with GraphPad Prism (version 10.0.0 for Windows, GraphPad Software, Boston, MA, USA).

The following measurements were analyzed with an ANOVA test with two factors systematically followed by a *post hoc* Tukey multiple comparison test:

- The cell proportions (dead, sonoporated, and PI+) and the eATP normalized signal (Fig 4 and 5) (pulse pressure and pulse length).
- The ATP is released per isolated cells and cells in responding-cell clusters (Fig 6A and 7A) (cell category and pulse length).
- The release rates (Fig 6C cell category and time; Fig 6B cell category and pulse length).
- The normalized model of release (Fig 7A) was analyzed first within the same ROI size, and then along time (ROI size and time).

**Fig 4.**
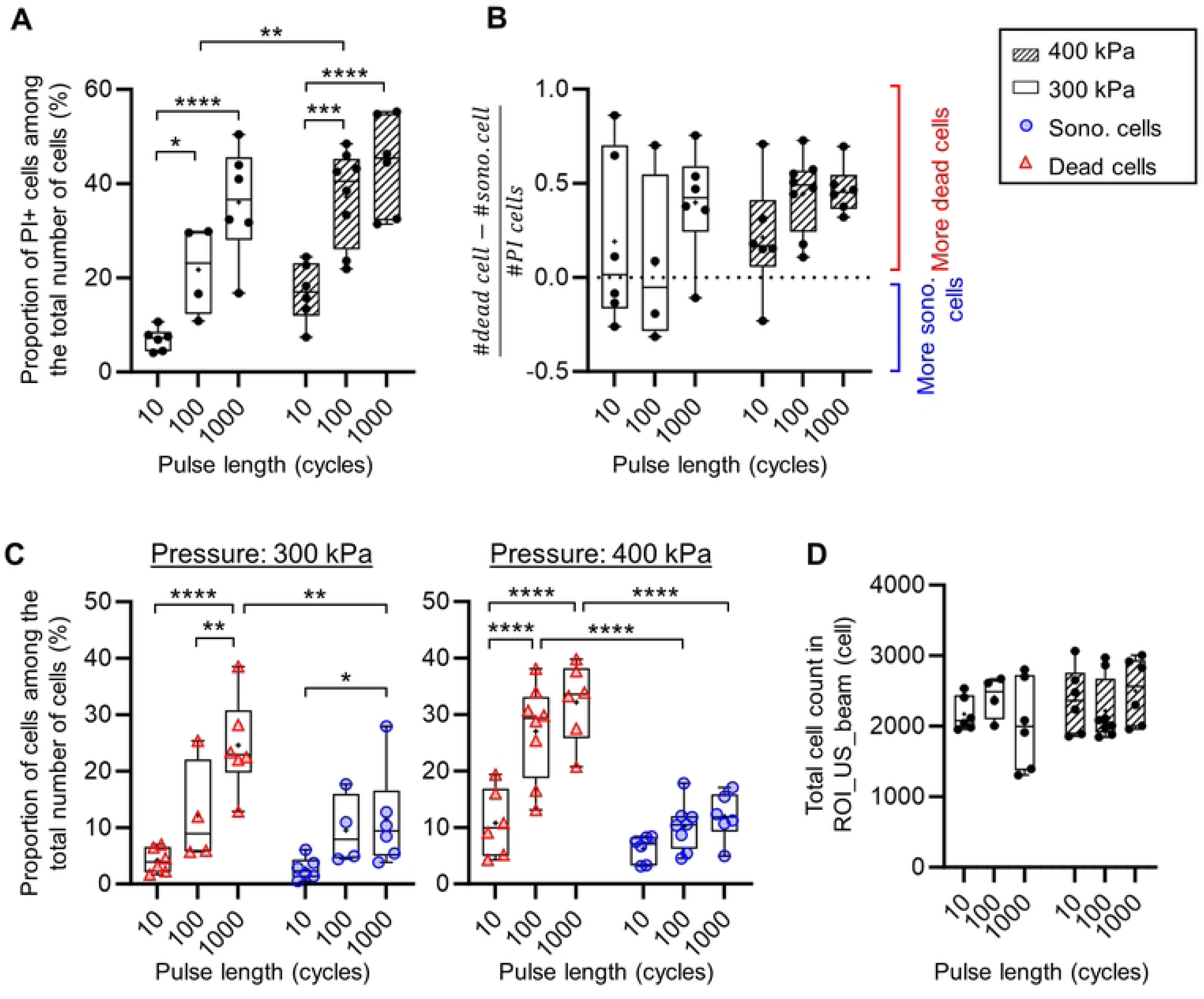
The PI-positive cell inside ROI_US_beam. (A) Graph of the proportions of PI-positive cells among the total number of cells counted inside ROI_US_beam in each chip. (B) Graph of the gap between the dead and the sonoporated cell counts inside ROI_US_beam in each chip showing predominance of dead or sonoporated cells. (C) Proportions of dead and sonoporated cells among the total number of cells counted inside ROI_US_beam in each chip. (D) Total cell count inside ROI_US_beam.

**Fig 5.**
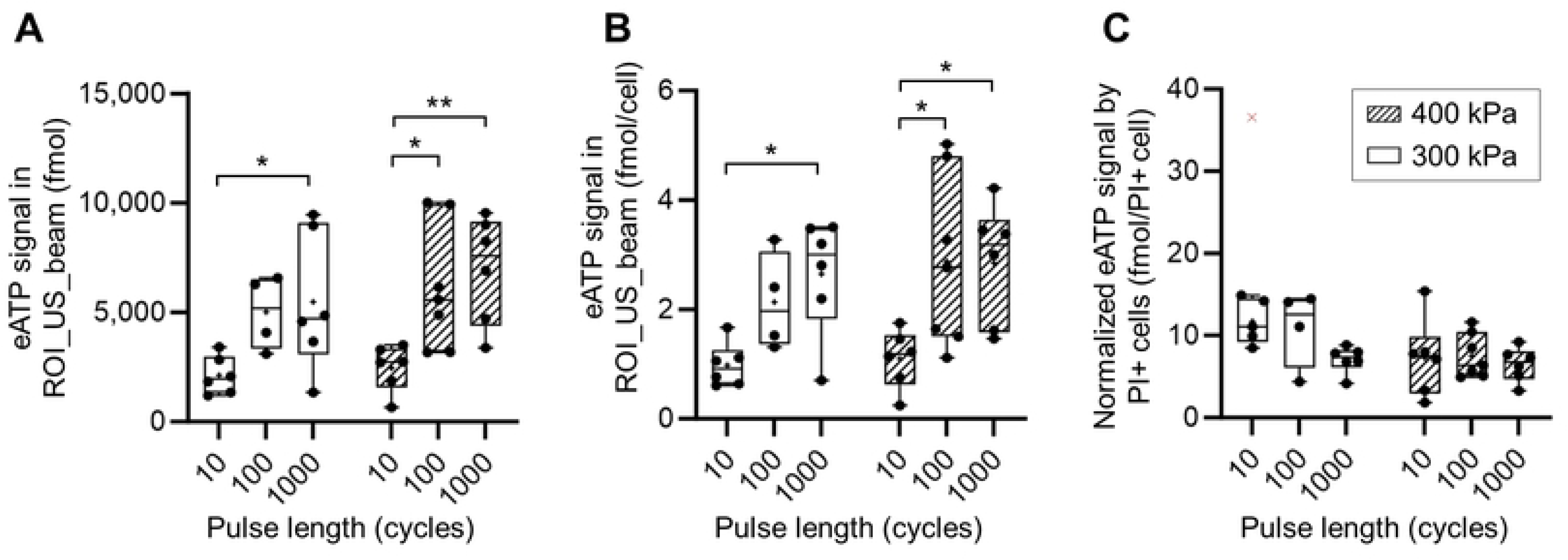
ATP released in the sonicated area. (A) Sum of the eATP signal inside ROI_US_beam. (B) Sum of the eATP signal inside ROI_US_beam normalized by the total number of cells in the area. (C) Sum of the eATP signal inside ROI_US_beam normalized by the cell count of PI-positive cells in the area.

**Fig 6.**
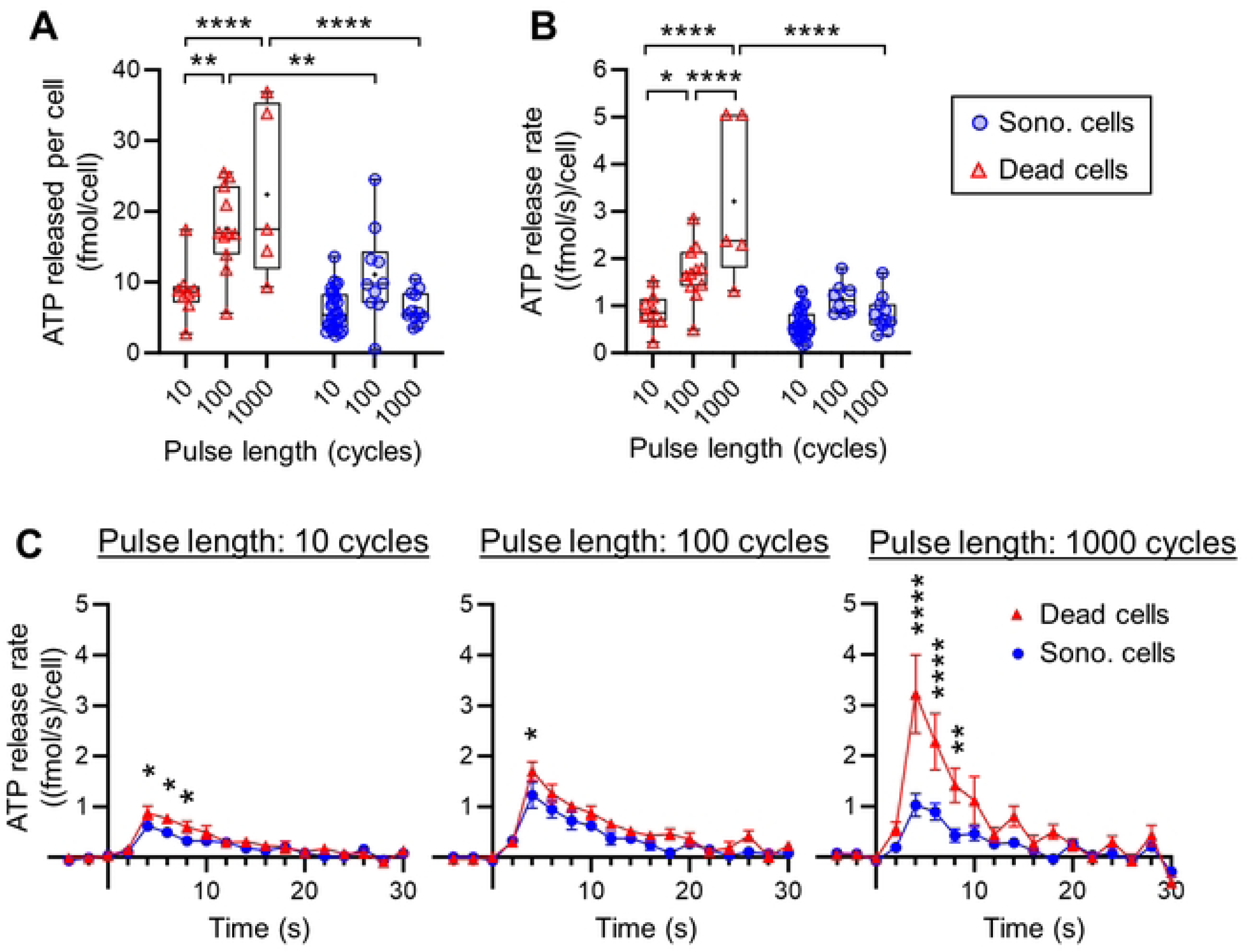
ATP release in isolated cells treated with 300 kPa. (A) ATP released by each isolated cell inside ROI_3.5px for each pulse group. 10 cycles pulse (cell analyzed: dead N = 8; sono. N = 28); 100 cycles pulse (cell analyzed: dead N = 11; sono. N = 10); 1000 cycles pulse (cell analyzed: dead N = 5; sono. N = 11). (B) Release rate calculated at 4 s for every pulse length for isolated cells inside ROI_3.5px. (C) Release rate (mean ± SEM) of ATP along time in isolated cells inside ROI_3.5px at different pulse lengths.

**Fig 7.**
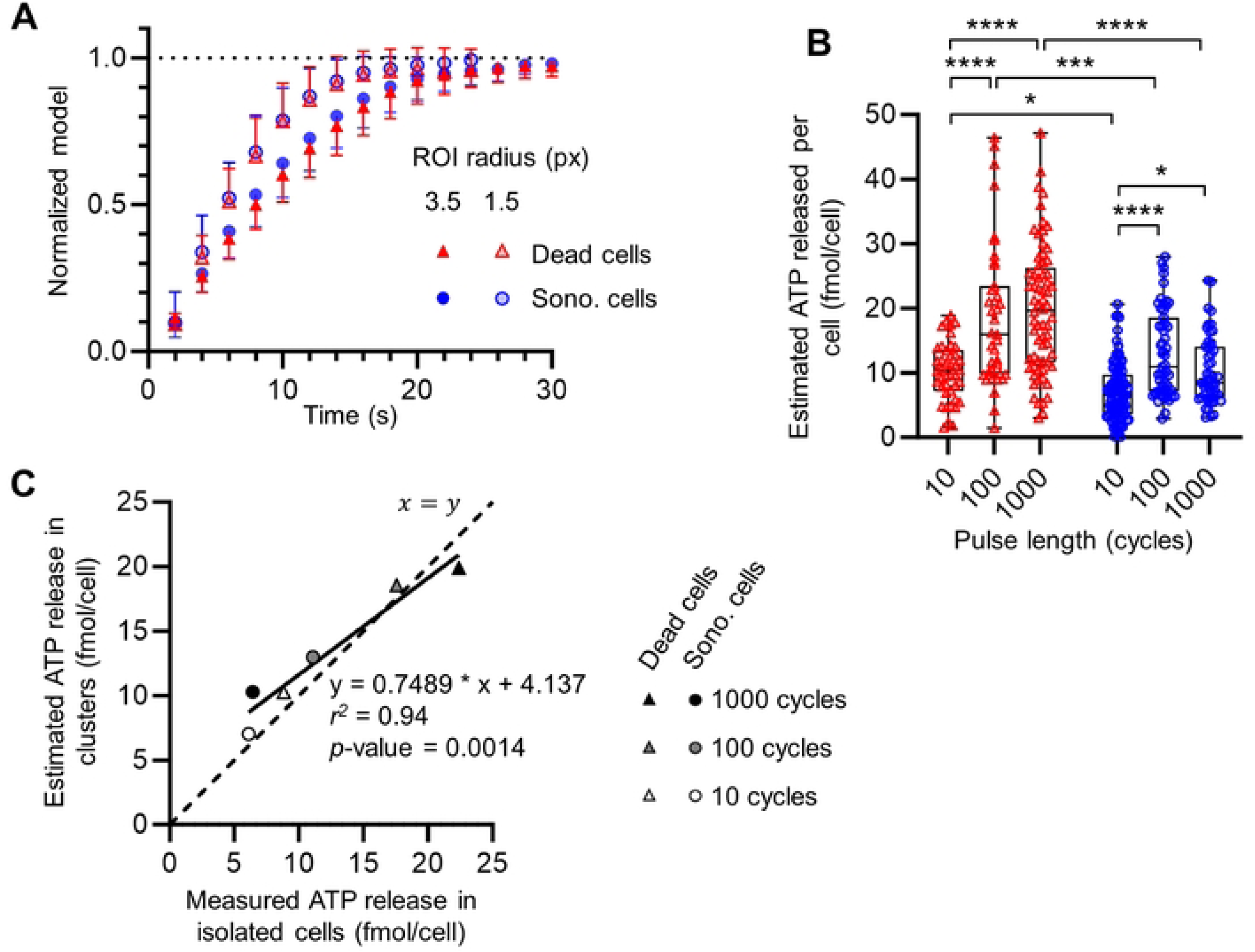
Estimation of ATP released in cells in responding-cell clusters. (A) Graph of the normalized models (number of model sono. = 24; dead = 49) of the first kinetic phase of release (mean and exterior SD) for measures inside ROI_3.5px in sonoporated cells (blue dots) and dead cells (red triangles) and inside ROI_1.5px in sonoporated cells (empty blue dots) and dead cells (empty red triangles). (B) Estimated ATP released by each cell inside the analyzed ROI_cluster using measurement of partial ATP released inside ROI_1.5px. (C) Graph of the average estimated ATP released using measurement of partial ATP released inside ROI_1.5px in ROI_cluster (calculated from the Fig 7B) as a function of the average measured ATP released in isolated cells (calculated from the Fig 6A).

The significance was indicated on graphs with ns. for *p* > 0.05, * for *p* < 0.05, ** for *p* < 0.01, *** for *p* < 0.001, and **** for *p* < 0.0001.

Linear regressions were used to analyze the relationship between the weighted and the detailed sum of eATP signal and the measurements in ROI_cluster (Fig 8).

**Fig 8.**
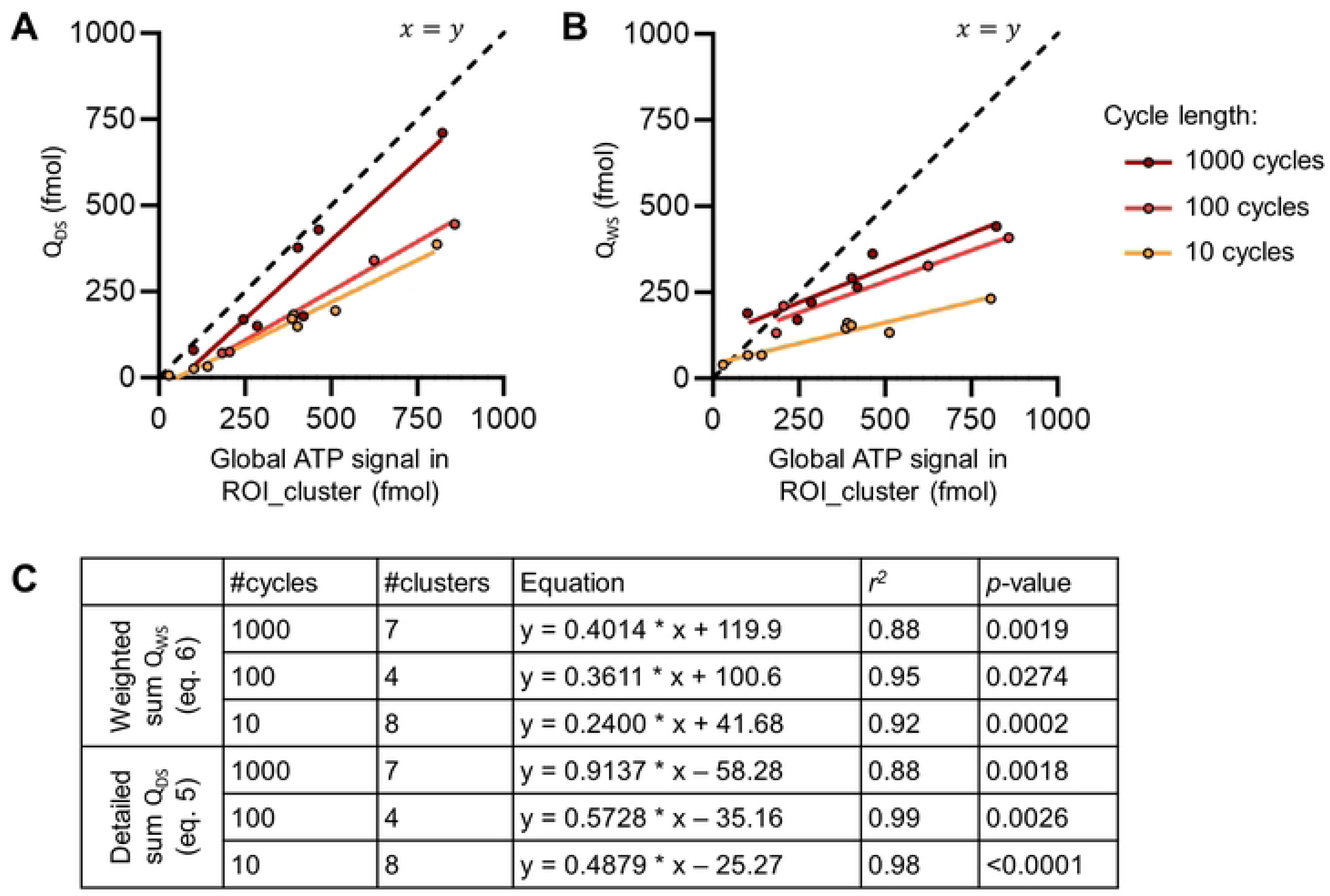
Correlation between global cell cluster measurement and individual signals. (A) Graph of the correlation between the detailed sum of ATP released by cells in clusters (eq. 5) with the global measurements done in the ROI_cluster corresponding. (B) Graph of the correlation between the weighted sum of ATP released by cells in clusters (eq. 6) with the global measurements done in the ROI_cluster corresponding. (C) Table of the fitted models, the coefficient of determination *r^2^*, and the associated *p*-values for the linear regressions.

Data were presented by pulse pressure separately to highlight the impact of the pulse length. Box plots, with a cross representing the mean, were used to describe the different measurements per pulse length. In the text, data are written in the format: mean ± standard deviation.

## Results and discussion

### Global impact of UTMC on cell poration status and ATP release

Within the tested conditions, a single UTMC pulse caused between 4% and 55% of PI+ cells in the ROI_US_beam (Fig 4A). This number increased with pressure and pulse length. Most chips had a greater count of dead cells (Fig 4B), particularly for the 400 kPa pulses. However, for the weaker pulses (300 kPa / 10 cycles and 300 kPa / 100 cycles) half of the chips presented a greater count of sonoporated cells, indicating, as expected, that milder US pulses have a gentler effect on cells.

The proportion of dead cells increased with the pulse length at both tested pressures (Fig 4C). However, the changes in the proportion of sonoporated cells were not significant except between the pulse 300 kPa / 10 cycles and 300 kPa / 1000 cycles. The dead cell proportion was significantly higher than the sonoporated cell proportion for stronger pulses i.e. 300 kPa / 1000 cycles, 400 kPa / 100 cycles, and 400 kPa / 1000 cycles. The average sonoporation proportion seemed to plateau around 10%, at 300 kPa between the pulse at 100 cycles and 1000 cycles (9.5 ± 6.2 % vs.11.5 ± 8.7 %; ns.) as well as for 400 kPa between the 100 cycles and the 1000 cycles (10.2 ± 4.2 % vs. 11.9 ± 4.2 %; ns.). The dead cell proportion seemed also to plateau in the pulse pressure 400 kPa between 100 cycles and 1000 cycles (27.1 ± 8.5 % vs. 32.2 ± 6.9 %; ns.). This observation of the plateau in dead cells and PI+ cell proportions at 400 kPa between 100 and 1000 cycles was probably caused by the finite number of MB present in the device.

Concomitantly, the global eATP measures inside ROI_US_beam (Fig 5A) also increased with the pulse length at both pressures, consistently with the fluorescence assessment of membrane poration. Thus, globally, the ATP release increased with PI uptake. The similarities of the global (Fig 5A) and the normalized (Fig 5B) ATP released showed the consistency in the cell density in the chips used. The normalization of the ATP release by the PI+ cell (Fig 5C) count showed that the overall average ATP released by PI+ cells was 8.2 ± 3.5 fmol. This value is in the range of the intracellular ATP quantity measured in the Triton assay detailed in SI (S5 Appendix).

### ATP release kinetics by isolated cells

To identify the contribution of sonoporated and dead cells to ATP release, we first analyzed the quantity of ATP released in isolated cells (Fig 6).

The analysis of the isolated cells was done on the 300-kPa-treated chips only because the 400-kPa-treated chips showed very few isolated cells (only 5 dead cells were found in one chip 400 kPa / 10 cycles). Looking through all our 300-kPa-treated chips, we found eATP signals in 72 individual cells in total, including 49 sonoporated cells (10 cycles: 28, 100 cycles: 10, 1000 cycles: 11 cells) and 24 dead cells (10 cycles: 8, 100 cycles: 11, 1000 cycles: 5 cells).

The ATP released was significantly higher in dead cells than in sonoporated ones for the 100 cycles (17.6 ± 5.9 vs. 11.1 ± 6.5 fmol/cell; *p* < 0.01) and 1000 cycles (22.4 ± 12.2 vs. 6.4 ± 2.3 fmol/cell; *p* < 0.0001) pulse (Fig 6A). The ATP released in the 10 cycles pulse was not significantly different between dead and sonoporated cells (8.8 ± 4.1 vs. 6.4 ± 2.3 fmol/cell; ns.). For sonoporated cells, the amount of eATP measured was not significantly impacted by the pulse length variation. Nevertheless, a slight increase of the ATP released was found with the 300 kPa / 100 cycles pulse which can be explained by the higher variability of measured values in this pulse. For dead cells, we found that the ATP released increased with the pulse length. The increase in ATP release with pulse length can be attributed to variations in either the pore size or opening duration. Indeed, the increase in pulse length is suggested to impact either the pore size or opening duration [19]. Then, longer pulses likely resulted in larger or longer-lasting pores, facilitating greater ATP release. Considering that depletion of ATP can induce cell death [39], at longer pulses, the larger or longer-lasting pores could allow more ATP to exit the cell leading to cell death. It was interesting to find that for the 10 cycles pulse, a similar ATP release was found for sonoporated or dead cells.

Surprisingly, the mean ATP released by isolated dead cells exceeded the intracellular ATP quantity estimated with the Triton assay (see S5 Appendix) in some conditions (up to 22.38 ± 12.24 fmol/cell for the pulse 300 kPa / 1000 cycles). Nevertheless, this amount of ATP released is still in the range found in the literature, where the intracellular ATP concentration is estimated to be up to 10 mM [40–42]. Therefore, considering the average estimated volume of our cells, this could allow a release of up to 100 fmol per cell. Additionally, it is worth noting that the intracellular ATP quantification in the Triton assay was performed with the luminometer on cells cultured in tissue culture-treated polystyrene dishes, whereas ATP release measurements were conducted with the EMCCD setup on cells cultured in PDMS chips. The use of detergent for ATP lysis, different apparatuses for the measurement and different substrates for cell growth could influence measurements.

As previously mentioned, these findings showed a systematic, linear-like decrease in the luminescence signal after the peak (Fig 2B). While sustained ATP release is typically observed *in vivo* [6,25,31], this pattern was absent in our current *in vitro* model. We hypothesize that this discrepancy may result, among other factors, from the limited number of cells in our model (approximately 10,000 cells per chip), which is well suited for analyzing individual cellular responses but does not fully recapitulate *in vivo* conditions. We cannot exclude the possibility that we could miss lower grade, and potentially longer lasting ATP release mechanisms mounting up to detectable levels *in vivo*.

The study of the release rate in isolated cells along time (ROI_3.5px) showed a similar pattern in the three pulses tested (Fig 6B). The release rate showed a peak at 4 s after the pulse and return to the baseline about 20 s after the pulse. The release time course of ATP and the peak of release rate were consistent with other studies using different ATP release stimuli, like mechanical stretching in a co-culture of AT1/AT2 cells [43] and in A549 [33,44]. Furthermore, our results revealed a faster release time course, and a release rate peak three orders of magnitude higher than those observed during non-lethal hypotonic shock [45]. This confirms that the release observed in our case likely involves a breach in the cell membrane. The findings showed a significantly higher release rate in dead cells compared to sonoporated cells between 4 s and 8 s for the 1000 cycles and the 10 cycles pulse and only at 4 s for the 100 cycles pulse (Fig 6C). This finding suggests that increasing the pulse length increased the chances of larger pores leading to cell death. As the 4 s time point after the pulse seems to be a determining time point, the statistical analysis was conducted at this time point. We found an increasing release rate with the pulse length in dead cells and a higher release rate in dead cells than in sonoporated cells at 1000 cycles (Fig 6B). This study of the release rate suggests that the amount of ATP released between the 2^nd^ and the 4^th^ second after the pulse is decisive for the future survival of the cell to pore formation.

### ATP released by cells inside responding-cell clusters

The isolated cell data offered a characterization of the ATP released by isolated cells. Nevertheless, we also wanted to analyze the release in responding cell clusters, i.e. eATP signals that quickly merged into a blob (Fig 3B & 3C).

To do so, we used the normalized models of the first kinetic phase of the isolated cells (eq. 4). Interestingly, we found that the averages of the normalized model at each time point showed very similar time course and release kinetics between sonoporated and dead cells (Fig 7A) with ROI_1.5px or ROI_3.5px. We can also observe that the curve of the average normalized model for ROI_1.5px was above the curve for ROI_3.5px, indicating that the ATP release in ROI_1.5px reached its vertex earlier than in ROI_3.5px. This could be explained by the diffusive transport of eATP outside ROI_1.5px before the end of the first phase of the release kinetic (Fig 2B). This confirmed our choice to use the ROI_3.5px to characterize isolated cell release. Moreover, at 4 s after the pulse, the fraction of eATP present inside ROI_1.5px (0.33 ± 0.11) was non-significantly different from the fraction of eATP inside ROI_3.5px (0.26 ± 0.06). As the fraction of ATP released inside the two ROIs at 4 s was not significantly different, we used an ROI of 1.5 pixels of radius and the associated fraction constant to estimate the total ATP released by cells inside cell clusters. The final ATP released by each cell inside the responding-cell clusters could be estimated by the amount of ATP released in ROI_1.5px at 4 s divided by 0.33, which is the fraction of ATP that was supposed to be already released at that time.

In total, 149 dead cells (10 cycles: 42, 100 cycles: 39, 1000 cycles: 68 cells) and 208 sonoporated cells (10 cycles: 102, 100 cycles: 55, 1000 cycles: 51 cells) were analyzed in 27 cell-responding clusters. The estimated ATP quantity released by cells inside the cell clusters (Fig 7B) was found to fall in the same range as the ones measured in isolated cells (Fig 6A). Moreover, the linear regression between the average estimated and measured ATP released quantity per cell (Fig 7C) showed an excellent determination coefficient (*r^2^* = 0.94) and matching of the average estimation with the measurements for all the pulse lengths for sonoporated and dead cells.

Considering the robustness of the estimated values, some precisions can be added to the conclusions drawn from isolated cell measurements based on fewer cells. First, the estimated released ATP confirmed a significantly higher release in dead cells compared to sonoporated cells for all pulse lengths tested. We could also observe the increase of the ATP released in dead cells between the 10 and 100 cycles pulses (10.3 ± 4.5 vs. 18.6 ± 11.2 fmol/cell; *p* < 0.0001) but we found the ATP released plateauing between the 100 and 1000 cycles pulses (18.6 ± 11.2 vs. 19.9 ± 9.9 fmol/cell; ns.). In sonoporated cells, we also found an increase between the 10 and 100 cycles pulses (7.0 ± 4.4 vs. 12.9 ± 6.7 fmol/cell; *p* < 0.0001) and a plateauing between the 100 and 1000 cycles pulses (12.9 ± 6.7 vs. 10.3 ± 5.3 fmol/cell; ns.) rather than a decrease as previously hypothesized in the analysis of isolated cells.

### Global validation of the cluster measurements

Single cell release based on the cluster analyses were validated by correlation analyses against the global ATP signal (ROI cluster) using two approaches.

The detailed sum (Q_DS_) was calculated with *q_i_*(eq. 5) representing the estimated ATP released by cell *i* at 4 s in ROI_1.5px described earlier. In this approach, a strong correlation was found between ROI_cluster and the Q_DS_ (determination coefficients *r^2^*: 0.98; 0.99; 0.88 for 10 cycles, 100 cycles, and 1000 cycles respectively) for all pulse lengths (Fig 8A). Interestingly, the best prediction using Q_DS_ was found for the 1000 cycles pulse (slope of 0.91). It is also the pulse where the dead cell count was the highest within the 300 kPa pulses (39.8% higher than the sonoporated cell count) (Fig 4C). Q_DS_ for 100 cycles and 10 cycles underestimated the total signal 64.4 ± 10.7 % and 54.6 ± 9.2 % respectively.

The weighted sum (Q_WS_) was also calculated, with *q_sc_*and *q_dc_* (eq. 6) representing the mean ATP released in isolated cells experiments (see Fig 6A). We used different *(q_sc_; q_dc_)* values for each pulse length were used to calculate Q_WS_ ((6.4, 22.4) for 1000 cycles; (11.1, 17.6) for 100 cycles; (6.1, 8.8) for 10 cycles). Q_WS_ was also positively correlated with the measurement in ROI_cluster (determination coefficients *r^2^*: 0.92; 0.95; 0.88 for 10 cycles, 100 cycles, and 1000 cycles respectively) (Fig 8B). Q_WS_ was also found to underestimate the measurements in ROI_cluster (14.1 ± 45.6 %, 31.4 ± 24.8 % and 47.2 ± 36.2 % for 1000 cycles, 100 cycles and 10 cycles respectively).

The substantial underestimation of the global ATP released in ROI_cluster with the sums in both cases can be explained by several factors. One factor that could partially explain the underestimation for Q_DS_ could be the underestimation of the number of cells by the sonoporation detection program, which detected ∼90% of PI+ cells based on the tested subsamples data set used for the code validation (see Fig D in S4 Appendix). Another possibility is that some cells could have detached from the substrate between the eATP signal acquisition and the viability assay which was done 40 min later. However, since the channels were coated with fibronectin (greater cell adherence) and since the total cell count after the pulse was not varying with the pulse length despite increasing dead cell counts (Fig 4D), this appears to be an unlikely explanation. Finally, the combined observation of a slight overestimation of the ATP released per cell inside cell-responding clusters (Fig 7C) with the observation of an underestimation of the global ATP released in ROI_cluster by both Q_DS_ and Q_WS_ could suggest the release of ATP by other pathways. Indeed, our image-based approach is more sensitive to acute bursts of ATP, such as expected with lytic release by pores. Indeed, eATP appears in the luminescence signal images as a punctual signal we use for the placement of a ROI. In our approach with ROI, we thus could miss slower release mechanisms, such as shear-activated [6] or eATP-driven ATP release [46] or other non-lytic release of ATP. With this reasoning, the measurements in ROI_cluster would encompass the signal of lytic and eventually non-lytic ATP release, whereas the sums only consider the release by lytic events. The observation of a closer prediction of the signals measured in ROI_cluster by Q_DS_ with the 1000 cycles pulse is supporting this hypothesis because of the high number of dead cells and predominance of lytic release.

### Impact of cell confluence on ATP release

The ATP released (Fig 9D) in LCD was higher than in HCD in sonoporated (LCD: 7.6 ± 4.1 fmol/cell vs. HCD: 4.6 ± 2.1 fmol/cell; *p* < 0.05) and dead cells (LCD: 12.9 ± 5.2 fmol/cell vs. HCD: 7.7 ± 2.8 fmol/cell; *p* < 0.001) as well. The decrease of the ATP quantity contained in cells with the increase of the cell density is consistent with the results obtained in the Triton assay (see S5 Appendix). We also found that the surface occupied (Fig 9D) by cells at LCD was bigger than the HCD (2425 ± 1188 µm^2^ vs. 1510 ± 543 µm^2^; *p* < 0.0001 Student’s *t*-test). The average volume of the cells was estimated at 7.55 ± 2.71 pL and 12.12 ± 6.36 pL at HCD and LCD, respectively. By considering that the intracellular ATP amount is at least the amount released during cell death, we arrived at an estimated intracellular ATP concentration of 1.06 ± 0.98 mM and 1.01 ± 0.73 mM of ATP. Our approach enabled us to conclude that the cells at low density released more ATP, likely because they contain higher ATP levels.

**Fig 9.**
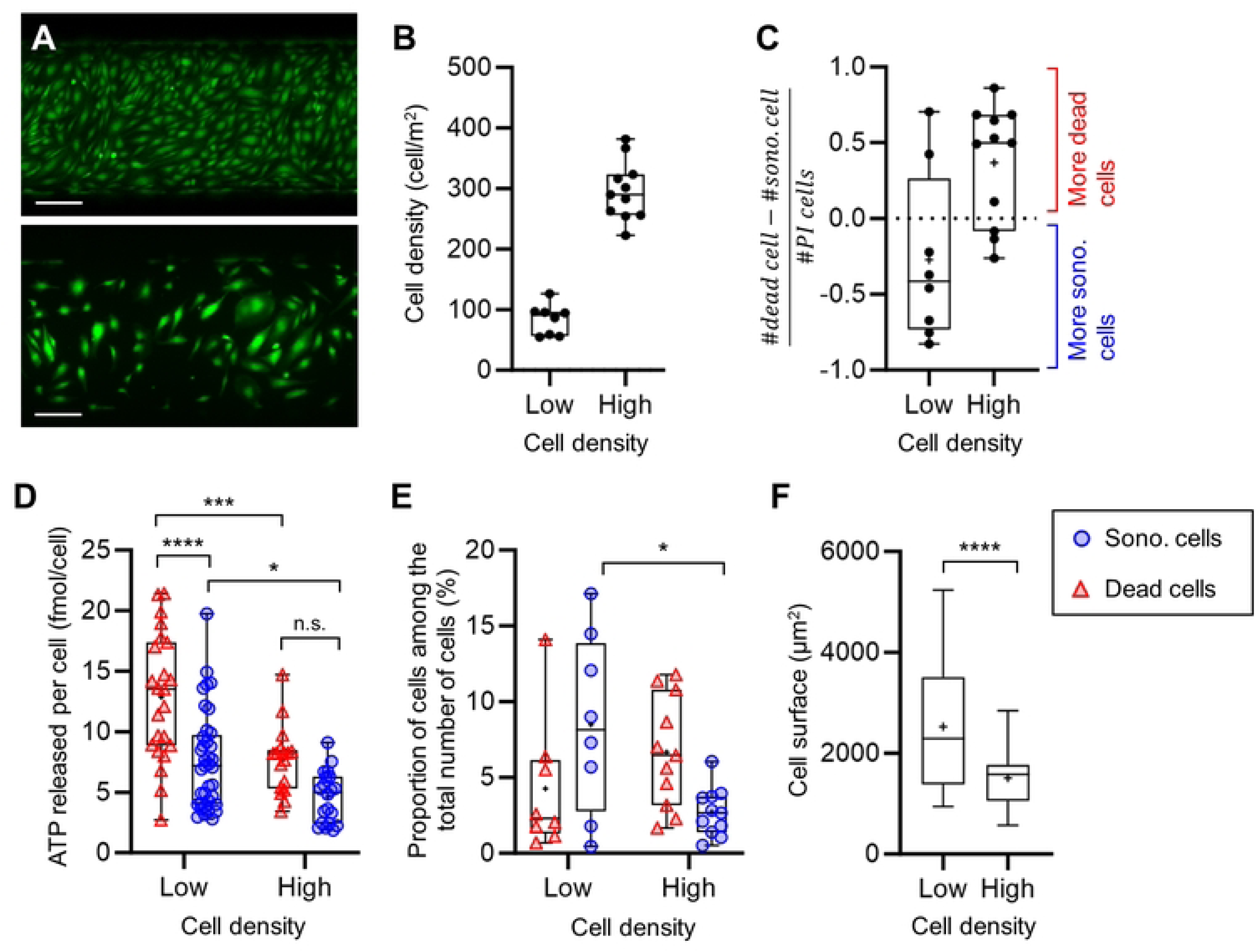
Impact of the cell density on the sonoporation and the ATP released. (A) Images of a region of two chips at high and low density. Scale bars: 200 µm. (B) Graph of the cell counts in the chips at low (N = 8 chips) and high (N = 11 chips) cell density. (C) Graph of the gap between the dead and the sonoporated cell counts inside ROI_US_beam in each chips. (D) ATP released per isolated cells at low (cell analyzed: dead N = 23; sono. N = 37) and high (cell analyzed: dead N = 17; sono. N = 20) cell density. (E) Dead and sonoporated cell proportions among the total number of cells in ROI_US_beam. (F) Cell surface of cells at low (N = 52) and high (N = 55) cell density.

The sonoporation proportion (Fig 9F) was higher at LCD (LCD: 8.5 ± 5.9 % vs. HCD: 2.7 ± 1.6 %; *p* < 0.05). We could see a higher proportion of dead cells in the high cell density group (HCD: 6.7 ± 3.6 % vs. LCD: 4.3 ± 4.5 %; ns.), but the results are not significant. Moreover, the results suggested that the chips with LCD presented more sonoporation than cell death (Fig 9C). These points support that cells at low density are more resilient to membrane poration. This could be explained by the fact that at low density, cells could contain more ATP due to their bigger size and could be able to sustain a higher amount of ATP depletion. As mentioned earlier, the pore size may vary with the US pulse length. Consequently, on a larger cell membrane, a given pore could represent a smaller breach, potentially easier to heal. At the same time, the MB concentration was the same in all the groups, making the MB/cell ratio higher at lower cell density which implies a higher chance of poration of a cell.

## Conclusions

Overall, we developed a microfluidic approach that allowed combining US and quantitative mapping microscopy to characterize the source and kinetics of ATP release following UTMC therapy *in vitro* in HUVEC. This calibrated and localized technique enabled us to quantify the ATP release by dead and sonoporated cells in femtomole quantities, consistently with eATP measurements reported in the literature, and which attests to the robustness of our general findings.

We found that dead cells are globally releasing more ATP than sonoporated cells, and have a higher release rate in the earliest time points after UTMC. These effects depended on the ultrasound pulse, and were consistent with the concept that stronger ultrasound (pressure and pulse length) caused larger pores in the cell membrane, eventually becoming unresealable, and thus leading to cell death. Interestingly, we found a similar ATP release kinetic profile for all cells, suggesting that the dominant release mechanisms were similar and likely involved pore formation in dead cells as well as sonoporated cells. In both cases, our findings support that, at the scale of a single cell, most, if not all, the ATP is released acutely in 10 to 50 seconds after the US pulse.

Our study supports that ATP can be released by endothelial cells even with brief, low-intensity UTMC pulses, allowing the potential to stimulate vasodilation and preserve the viability of endothelium. Understanding the release mechanisms and the reservoirs of UTMC-mediated ATP release provides valuable insights for enhancing radiotherapy, particularly through the potential of UTMC as a radiosensitizer in tumors.

## Acknowledgment

The authors thank the CRCHUM Microfluidics Core Facility supported by the TransMedTech Institute and its main financial partner, the Canada First Research Excellence Fund, and specifically Dr. Amélie St-Georges-Robillard for technical support during the microfluidic design process and manufacturing. We also thank Boris Chayer, M. Sc. research engineer at the laboratory, for technical support.

## Supporting information captions

S1 Appendix. Transducer calibration and characterization

S2 Appendix. Microfluidic device manufacturing

S3 Appendix. Calibration factor for eATP quantification

S4 Appendix. Cell classification program description and validation

S5 Appendix. Estimation of total ATP in HUVEC cells using an independent method (Triton assay)

